# Tissue-specific 3 prime-end adenylation of miR-125b mediates cell survival

**DOI:** 10.1101/672436

**Authors:** Matthew T. Blahna, Jean-Charles Neel, Ge Yu, Justin Louie, Prajakta Ghatpande, Giorgio Lagna, Akiko Hata

## Abstract

Next-generation sequencing has uncovered microRNAs (miRNAs) that undergo sequence modifications, known as isomiRs. Their physiological significance, however, remains uncertain, partly because they generally comprise a small fraction of total miRNAs. Here we report that more than 60% of miR-125b, one of the most abundant miRNAs in vascular smooth muscle cells (vSMC), exists as an edited isoform containing a non-templated adenosine residue at the 3 prime-end (miR-125b+A). The properties of miR-125b+A, such as stability and subcellular localization, are similar to those of canonical miR-125b, but miR-125b+A more potently inhibits the expression of a subgroup of targets, including the apoptosis effector Caspase-6 (*CASP6*). In the *CASP6* transcript, adenylated miR125b preferentially targets a conserved, atypical site, with an unusual 36 nucleotides loop between seed sequence and 3 prime-end supplementary site. PAP associated domain containing 2 (PAPD2) is responsible for monoadenylation of miR-125b. Downregulation of PAPD2 results in the conversion of miR-125b+A to miR-125b, derepression of *CASP6,* and sensitization of vSMC to apoptotic stimuli. Thus, atypical site recognition by a tissue-specific isomiR fulfills a pro-survival role.

## Introduction

MiRNAs are small noncoding RNAs, approximately 22-nucleotide (nt) long that generally post-transcriptionally repress gene expression ^1,2^. Dysregulation of miRNAs accounts for various pathological or physiological processes and developmental abnormalities ^3,4^. More than 2,500 miRNAs have been thus far annotated in humans ^5^. The canonical miRNA biogenesis pathway begins with transcription of the miRNA gene by RNA polymerase II. This transcript, named “pri-miRNA”, is processed by the RNase III enzyme Drosha to generate a hairpin-structured “pre-miRNA” in the nucleus, which is then processed by the enzyme Dicer in the cytoplasm to produce the mature miRNA^2^. Next-generation sequencing of small RNA libraries has identified sequence variations in both pre-miRNAs and mature miRNAs ^6-11^. These sequence variants, collectively known as “isomiRs”, include either nucleotide substitutions and addition or deletion of mono- or oligo-nucleotides to the 5’- or 3’-end. isomiRs with additional nucleotide(s) result from differential processing (genomically encoded or templated) or post-processing modification by nucleotidyltransferase (nontemplated), or a combination of both ^6-11^. The general abundance of adenosine (A) or uridine (U) nucleotides added at the 3’-end of miRNAs is constant across invertebrates and vertebrates^11^, suggesting that miRNA tailing is evolutionally conserved^10,11^. Furthermore, 3’-adenylation of several miRNAs is dynamically regulated throughout development in *Drosophila* ^12^ and aberrant expression of specific isomiRs has been reported in cancer cells ^8,13^.

Several enzymes can mediate nontemplated nucleotidyl additions to the 3’-end of miRNAs. For example, terminal uridylyl transferase TUT4 (also known as TUTase4, ZCCHC11 or PAPD3) and TUT7 (also known as PAPD6) are known to induce oligouridylation of pre-let-7, which blocks Dicer processing and facilitates miRNA decay by the 3’-to-5’ exonuclease DIS3L2^14-19^. TUT4 also mediates the 3’-uridylation of miR-26 and modulates its silencing activity without altering the stability of the miRNA ^20^. PAP associated domain containing 4 (PAPD4, also known as GLD2) promotes the 3’-monoadenylation of liver-specific miR-122, which increases its stability ^21,22^. Conversely, 3’-monoadenylation of a 23-nt-long isomiR of miR-21 (miR-21+C) by PAPD5 (also known as TUTase3 or TRF4-2) leads to degradation ^13^. Finally, a poxvirus-encoded poly (A) polymerase mediates adenylation and degradation of host miRNAs ^23^.

In addition to the effects on miRNA stability and silencing activity, it has been suggested that miRNA tailing plays a role in the selective affinity between miRNAs and different Argonaute proteins ^24^ or in the translocation of isomiRs to the nucleus and sequestration from target mRNAs ^25^. However, the physiological significance of isomiR generation remains unknown, partially because they are often found in limited abundance (less than 5% of total miRNA population).

In this study, we report the existence of an isomiR of miR-125b that is enriched and abundantly expressed in PASMC. We find that 3’-monoadenylated miR-125b (miR-125b+A) is the most abundant form of miR-125b in PASMC, being expressed at a ∼2:1 ratio to the unmodified form (miR-125b+A: miR-125b). Tailing of miR-125b+A requires the nucleotidyl transferase PAP associated domain-containing 2 (PAPD2, also known as TUTase6, TUT1 or Star-PAP). Reversion of miR-125b+A to miR-125b sensitizes PASMC to apoptotic stimuli because it reduces the silencing of pro-apoptotic Caspase-6 (CASP6), while it increases the silencing of IL-1β transcripts. Furthermore, targeting of *CASP6* by miR125b+A involves a 3’ supplementary site different from that targeted by canonical miR125b. This study provides evidence of the fine-tuning of miRNA targeting by highly efficient, cell type-specific, nontemplated tailing, resulting in gene expression changes that promote cell survival.

## Results

### High rate of 3’-monoadenylation of miR-125b in PASMC

To identify miRNAs involved in the regulation of vSMC biology, we performed “next generation” sequencing of small RNAs (miR-Seq) using triplicate RNA samples from human primary pulmonary arterial smooth muscle cells (PASMC) untreated or treated with BMP4 for 24 h (Fig. 1A). Alignment of sequencing reads to the miRBase database (Release 19.0, http://www.mirbase.org/) confirmed that miR-10a, miR-21, miR-100 and miR-143-3p, all known to modulate SMC differentiation ^26,27^, were among the most abundant miRNAs in PASMC, regardless of BMP4 stimulation (Fig. 1A). Similarly, miR-125b-5p is also a key regulator of SMC function ^28-30^ and was highly abundant (Fig. 1A).

**Fig. 1:**
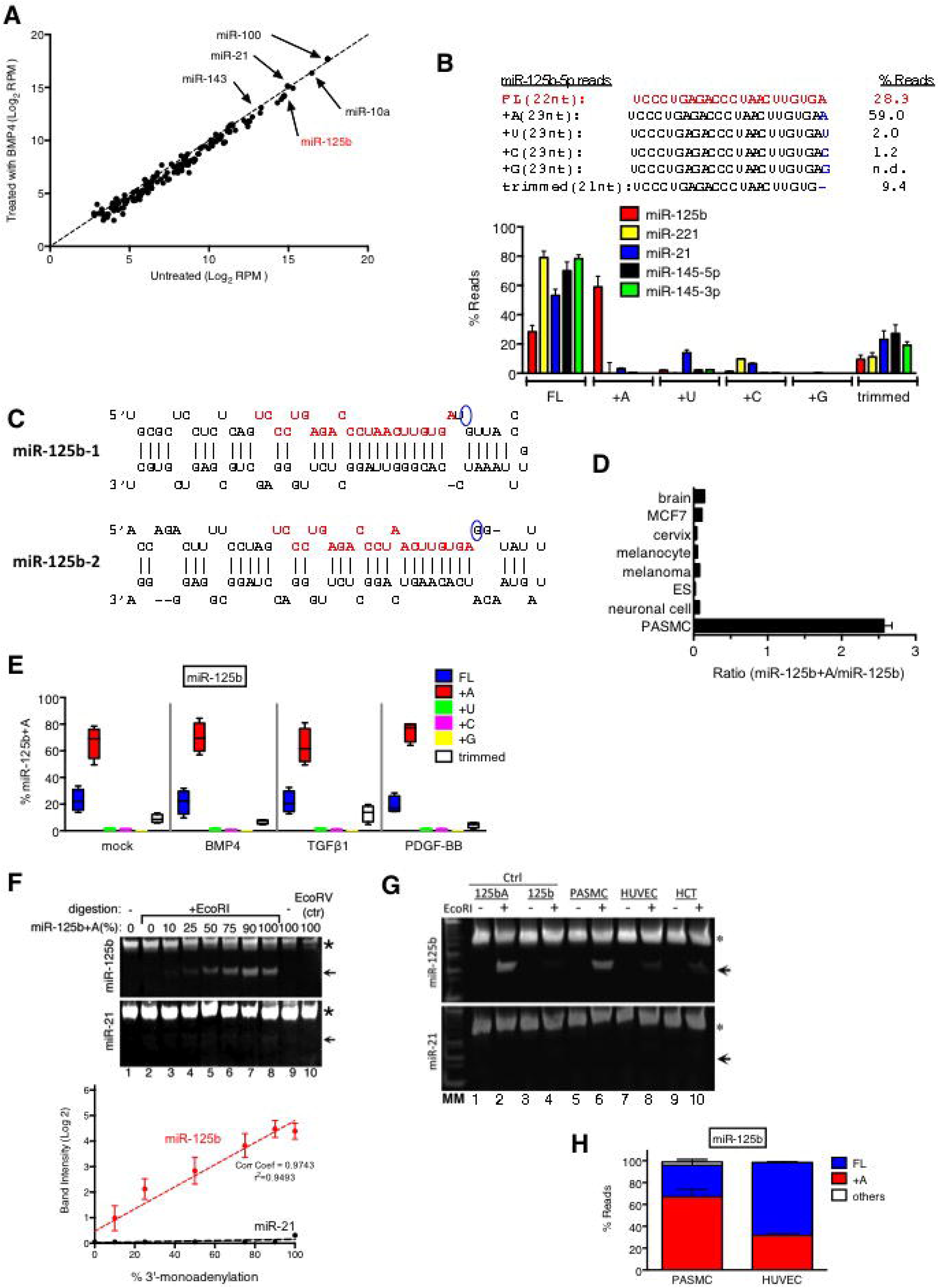
Large fraction of 3’-monoadenylated miR-125b found in PASMC. **A.** PASMC unstimulated (control) or stimulated with BMP4 for 24 h prior to miR-Seq analysis. Amounts in reads per million (RPM) of different miRNAs expressed in PASMC in BMP4 treated cells (Y axis) were plotted against untreated cells (X axis) in Log2. Five abundant miRNAs in PASMC (miR-100, -10a, -125b, -21, -143-3p) are indicated. **B.** Heterogeneity of miR-125b sequences found in PASMC. Sequence of canonical 22 nt miR-125b sequence (FL, red), miR-125b isomiRs that include a single nucleotide added at the 3’-end (+A, +U, +C, or +G) or one nucleotide shorter than canonical miR-125b (trimmed, 21 nt) and average percentage of those sequence in three independent miR-Seq libraries from PASMC are shown. **C.** The pre-miRNA structures of the two miR-125b genes in the human genome are shown. The red sequences indicate mature miR-125b, and the blue circles the nucleotides adjacent to the 3’-end of the miR-125 sequence. **D.** Mean ± SEM ratio of miR-125b+A reads to total miR-125b reads from small RNA sequencing libraries derived from various cancer (melanoma, breast carcinoma MCF7) or normal tissues (ES, neuronal cells, melanocyte, cervix, brain), based on miR-Seq data available at YM500v2 and miRBase and our result in PASMC. **E.** Abundance of miR-125b isomiRs in PASMC is not altered upon growth factor stimulation. Average fraction (%) of miR-125b (FL) or other 3’-isomiRs (+A, +U, +C, +G, or Trimmed) reads compared to total miR-125b reads in PASMC untreated (mock) or treated with BMP4, TGF-β1 or PDGF-BB for 24 h. Results are shown as mean±SEM of three independent libraries. **F.** Cos7 cells transfected with different ratios of miR-125b and miR-125b+A mimic were subjected to the LD assay using primers to detect miR-125b or miR-21 (control). The samples are; lane 1, untreated; lanes 2-9, digested with EcoRI; and lane 10, treated with EcoRV as negative control. Intensities of EcoRI-digested bands (indicated with an arrow) from top (miR-125b, red) or bottom panels (miR-21, black) were plotted against the calculated ratio (%) of miR-125b+A vs miR-125b. Cos7 cells transfected with known miRNA-125b sequences were used as a positive control. Asterisk indicates non-specific fragments. **G.** The LD assay was applied to RNA samples derived from PASMC (lanes 5-6), HUVEC (lanes 7-8), or HCT116 cells (lanes 9-10). RNA samples from Cos7 cells transfected with miR-125b+A (125bA; lane 1-2) or miR-125b (125b; lanes 3-4) were used as controls. MM: molecular markers. For each RNA sample, odd lanes are untreated and even lanes are EcoRI-treated samples. **H.** Small RNA libraries derived from PASMC and HUVEC were subjected to miR-Seq. The abundance of reads of canonical miR-125b (FL), miR-125b+A (+A) or other miR-125b isomiRs (others) were calculated as percentage of total miR-125b reads and plotted. Results are shown as mean±SEM of two (HUVEC) or four (PASMC) independent libraries.

However, only 28.2±4.5% of the miR-125b-5p sequence reads matched the 22-nt full-length sequence (5’-UCCCUGAGACCCUAACUUGUGA-3’), to which we refer as “FL” or “miR-125b” (Fig. 1B, FL). Instead, most miR-125b-5p reads (59.0±7.6%) were 23-nt long and contained an additional adenosine at the 3’-terminus (Fig. 1B, +A). This variant of miR125b-5p, to which we refer as “miR-125b+A”, has not been reported previously, nor have other miR-125b isomiRs. Notably, miR-125b sequences with additional U, C or G nucleotides amounted to less than 3.2% of the total miR-125b sequences in our miR-Seq library, and a similarly low frequency of miRNA 3’-end modifications (tailings) was observed in different miR-Seq libraries (Fig. 1B)^11^. An analysis of the rate of 3’ tailing of the 24 most abundant miRNAs revealed a small fraction of nontemplated 3’-adenylation of miR-100 (30%) and miR-99a (29%) and possibly templated 3’-adenylation of miR-151a (15%), while other miRNAs exibited background levels (<10%) of adenylated isomiRs (Suppl. Fig. S1). Thus, a high abundance of 3’-monoadenylation is unique to miR-125b.

To examine the source of the large fraction of miR-125b+A, we first considered the hypothesis that miR-125b+A might be an alternative transcription product and referred to the two genomic loci that encode miR-125b in vertebrates: miR-125b-1 and miR-125b-2 (Fig. 1C) (miRBase). Sequences of both loci revealed that the nucleotides following the 3’-end of mature miR-125b-5p (5’-UCCCUGAGACCCUAACUUGUGA-3’, Fig. 1C, shown in red) in miR-125b-1 and miR-125b-2 are U and G, respectively (Fig. 1C, blue circles). It is highly unlikely that the abundance of miR-125b+A might be due to a single nucleotide polymorphism (SNP) of these genomically encoded nucleotides, in conjunction with a Dicer processing error leaving one additional nucleotide at the 3’ end, as such substitutions are not found in the NHLBI GO Exome Sequencing Project (ESP) database (http://evs.gs.washington.edu/EVS/). Furthermore, the preponderance of the miR-125b+A isoform is unique to PASMC as the ratio (*r*) of miR-125b+A to total miR-125b reads in different tissue samples that express miR-125b is *r* <0.2, according to data from the YM500v2 miR-Seq database (http://ngs.ym.edu.tw/ym500v2/index.php) and miRBase, while on average *r*=2.5 in PASMC (Fig. 1D). The fraction of miR-125b+A isoform was unaltered by the treatment with different growth factors that affect PASMC phenotype, such as BMP4, TGFβ and PDGF-BB (Fig. 1E). We also compared the relative abundance of miR-125b+A in PASMC lines established from five unrelated individuals and found no significant difference among subjects (data not shown). Therefore, the high abundance of the miR-125b+A isoform is an intrinsic characteristic of PASMC and likely to be a product of a post-transcriptional modification of canonical miR-125b.

As conventional qRT-PCR methods to monitor miRNA abundance do not accurately quantitate the amount of miR-125b+A when both miR-125b and miR-125b+A are expressed in the same tissue, we developed a novel semi-quantitative method, the “Ligation/Digestion (LD) Assay”, to measure the two miR-125b isoforms. Briefly, an RNA adaptor whose sequence starts with 5’-UUC-3’ was ligated to the 3’-end of small RNAs to create a cDNA library. When the adaptor is ligated to RNA molecules ending with the 5’-GA<u>A</u>-3’ sequence, such as miR-125b+A but not canonical miR-125b, it generates an EcoRI recognition site (5’-GAATTC-3’) upon conversion to cDNA by reverse transcriptase (RT). A specific miRNA was amplified by PCR using a miRNA sequence-specific forward primer and a reverse primer that anneals to the adaptor (Suppl. Fig. S2). The PCR products were digested with EcoRI and the resulting fragments were separated by agarose gel electrophoresis. To test and titrate the system, different ratios of miR-125b and miR-125b+A synthetic RNA oligonucleotide mixtures were transfected into Cos7 cells, which express low amounts of endogenous miR-125. In cells left untransfected (Fig. 1F, lane 1) or transfected only with miR-125b (Fig. 1F, lane 2), no EcoRI digested band appeared (Fig. 1F, top panel), indicating that nonadenylated miR-125b cannot be detected. Other miRNAs whose sequence ends with 5’-GAA-3’ are not detected with a forward primer specific for miR-125b. No digested fragments were detected when samples were not treated with EcoRI (Fig. 1F, lane 9) or treated with EcoRV (Fig. 1F, lane 10), confirming that the digestion is dependent on the EcoRI recognition sequence generated as a result of 3’-monoadenylation of miR-125b. miR-21, which is endogenously expressed in Cos7 cells, has a sequence ending in 5’-GA-3’, identical to non-adenylated miR-125b, but we have not observed any 3’-monoadenylated miR-21 isomiR in our miR-Seq data. Unsurprisingly, we did not detect an EcoRI digested band in the LD assay using a specific primer for miR-21 (Fig. 1F, bottom panel), confirming that the assay detects specifically only 3’-monoadenylated miRNAs ending in 5’-GA-3’. When increasing ratios of miR-125b+A over miR-125b were transfected into Cos7 cells (keeping constant the total transfected miRNA amount), we observed an increase of the digested fragments in linear correlation with the abundance of miR-125b+A between 25%-80% (Fig. 1F, red line, bottom panel). Using these data, we estimated the abundance of miR-125b+A in PASMC, human umbilical cord vein endothelial cells (HUVEC) and human colon carcinoma HCT116 cells at about 60-70% for PASMC (Fig. 1G), which is consistent with the result of miR-Seq (Fig. 1D), and 20-30% in HUVEC and HCT116 (Fig. 1G). We confirmed the abundance of miR-125b+A in HUVEC by miR-Seq, which yielded a fraction of 31% miR-125b+A/total miR-125b in HUVEC compared to ∼60% in PASMC (Fig. 1H). These results confirm that the high abundance of miR-125b+A is PASMC-specific.

### PAPD2 is required for 3’-monoadenylation of miR-125b in PASMC

To identify an adenyltransferase responsible for 3’-monoadenylation of miR-125b in PASMC, we performed an RNAi screen of different noncanonical poly(A) polymerases endowed with adenyltrasferase activity (Fig. 2A) and measured the amount of miR-125b+A by the LD assay (Fig. 2B). The knockdown of PAPD2 by siRNA resulted in a decrease of miR-125b+A, while no change was observed by knockdown of PAPD1, 4, 5, or 7, despite similar levels of suppression of all PAPDs upon siRNA transfection (Fig. 2B, right panel, qRT-PCR). It is also notable that the total amount of all forms of miR-125b, as measured by qRT-PCR, was not altered by si-PAPD2 treatment (Fig. 2B, right panel, red bars), suggesting that there is no significant change of the stability of miR-125b upon tailing. We also confirmed the role of a subgroup of PAPDs in PASMC and HUVEC by miR-Seq. PASMC transfected with si-Control, si-PAPD2, si-PAPD4, or si-PAPD5 were subjected to miR-Seq to measure the number of miR-125b and miR-125b+A reads (Fig. 2C). While knockdown of PAPD4 or PAPD5 had no detectable effect, PASMC depleted of PAPD2 (siPAPD2-PASMC) displayed a significant decrease of the miR-125b+A fraction from 55% to 35.5% (Fig. 2C, miR-125b). The rate of adenylation of miR-100 was not affected in siPAPD2-PASMC, suggesting that PAPD2 is responsible specifically for adenylation of miR-125b (Fig. 2C, miR100). Surprisingly, the fraction miR-125b+A found in HUVEC (∼24%) was unchanged upon PAPD2 downregulation (Fig. 2C, HUVEC), suggesting that the small fraction of miR-125b+A in HUVEC is generated by an adenyltransferase different from PAPD2. Notably, the fraction of miR-151a bearing an additional A at the 3’ was decreased upon knockdown of PAPD2 in both PASMC (from 24% to 10%) and HUVEC (from 29% to 5%). This result suggests that miR-151a might be adenylated by PAPD2, similarly to miR-125b, despite the additional A being potentially encoded in the genome. These results demonstrate a highly tissue-specific and efficient substrate-enzyme relationship between miR-125b and PAPD2 in PASMC and suggest the involvement of a PASMC-specific mechanism that enhances the specificity and efficacy of monoadenylation of miR-125b by PAPD2 to produce a large fraction of miR-125b+A in PASMC.

**Fig. 2.**
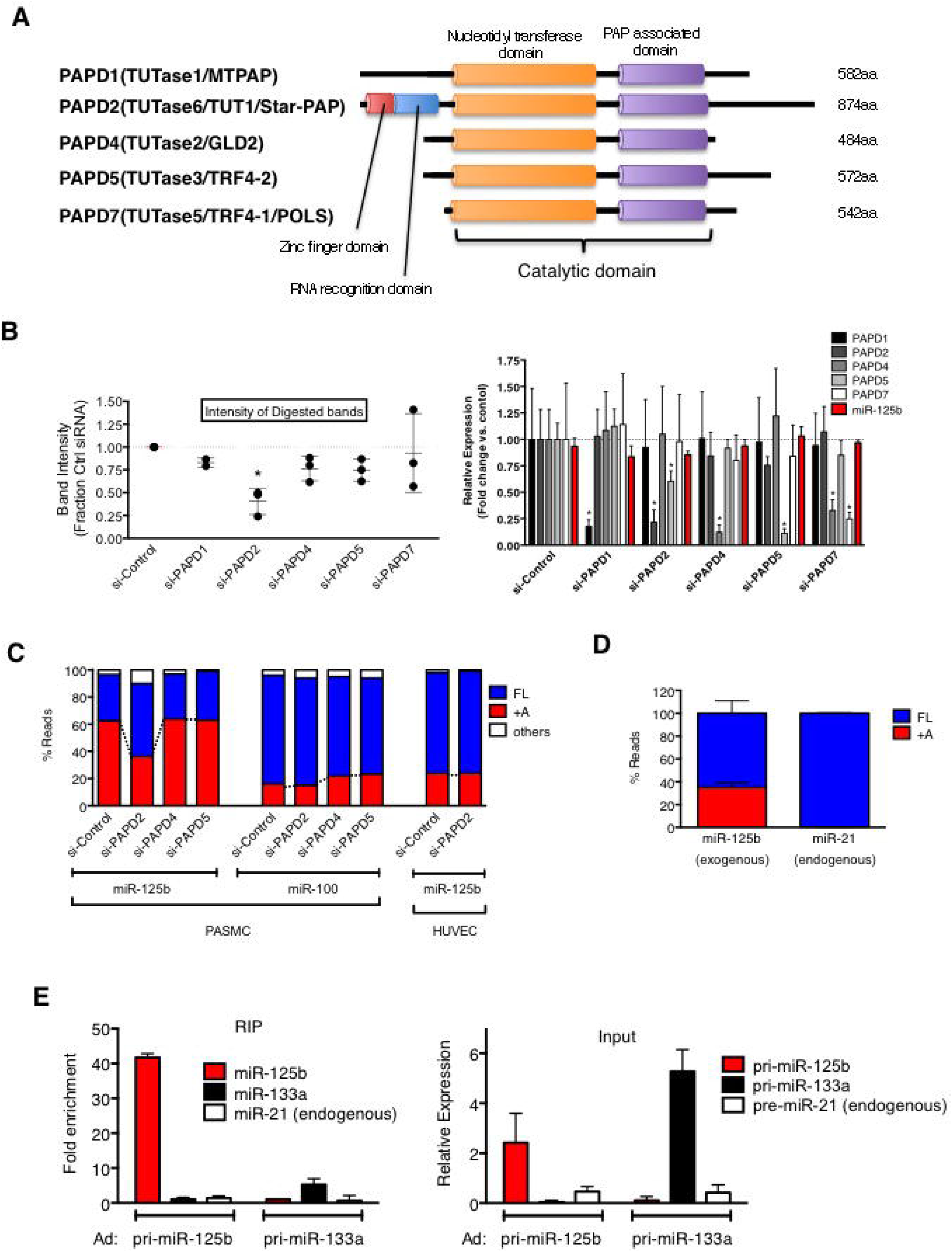
PAPD2 is required for the appearance of miR-125b+A. **A.** Schematic representation of domain structures of five members of the PAPD family with adenyltransferase activity. The right column states the total amino acid (aa) length of each protein. **B.** Application of the LD assay to RNA samples from PASMC transfected with siRNAs against PAPD1, PAPD2, PAPD4, PAPD5, PAPD7 or control. 3’-monoadenylation of miR-125b was examined. The digested bands were quantitated and plotted as relative intensity normalized to si-Control (left panel). Results are shown as mean±SEM of three independent assays. Efficiency and specificity of downregulation of different PAPDs by siRNAs were examined by qRT-PCR analysis of PAPD1, 2, 4, 5, and 7 mRNAs and miR-125b (right panels). Results are shown as mean±SEM of three independent samples identical to the samples used for the LD assay. **C.** Small RNA libraries were generated with RNA samples from PASMC or HUVEC transfected with siRNAs against PAPD2, PAPD4, or PAPD5 or control and subjected to miR-Seq. Abundance of canonical (FL), monoadenylated (+A), or other 3’-tailing (others) of miR-125b or miR-100 were calculated as percentage of read counts of each sequence divided by total miR-125b or miR-100 read counts. **D.** Small RNA libraries were generated from Cos7 cells expressing adenovirus-transduced rat pri-miR-125b, followed by miR-Seq. Abundance of canonical and 3’-monoadenylated miR-125b and miR-21 reads were calculated as percentage of total miR-125b reads and plotted. Results are shown as mean±SEM of three independent libraries. **E.** RIP assay was performed to detect the interaction between adenovirus-transduced rat pri-miR-125b or pri-miR-133a and Flag-tagged PAPD2 using anti-Flag antibodies or control IgG. Results (RIP, left panel) are shown as fold enrichment of the amount of pre-miR-125b, pre-miR-133a, or pre-miR-21 in PAPD2 immunoprecipitates over IgG control. Levels of expression of pri-miR-125b and pri-miR-133a as well as pri-miR-21 (endogenous) were measured by qRT-PCR prior to immunoprecipitation (input, right panel). Results are plotted as relative levels of miRNA compared to uninfected cells. Mean±SEM of three independent experiments are shown.

To examine whether exogenously expressed primary transcripts of miR-125b (pri-miR-125b) can be monoadenylated, we transduced a rat pri-miR-125b adenovirus expression construct into Cos7 cells and performed a miR-Seq analysis. About 35% of exogenously expressed pri-miR-125b was processed to generate miR-125b+A in Cos7 cells (Fig. 2D). The rate of 3’-monoadenylation of exogenous miR-125b in Cos7 cells was lower than that of endogenous miR-125b in PASMC but comparable to that observed in HUVEC (Fig. 1H). Endogenous miR-21, which is highly expressed in Cos7 cells, was not monoadenylated (Fig. 2D). Notably, mature miR-125b expressed in Cos7 cells was not adenylated (Fig. 1F and 1G), suggesting that monoadenylation of miR-125b might be coupled with processing of pre-miR-125b to miR-125b by Dicer. Thus, we hypothesized that PAPD2 might associate with pre-miR-125b prior to Dicer processing. To test the interaction between PAPD2 and pre-miR-125b, we performed a UV crosslinking and RNA immunoprecipitation (RIP) assay (Fig. 2E). Cos7 cells were transduced with Flag-tagged PAPD2 and pri-miR-125b or pri-miR-133a (control). An anti-Flag antibody was used to pull down PAPD2, and in the precipitates we measured by qRT-PCR analysis the amount of pre-miR-125b, pre-miR-133a (exogenous control) or pre-miR-21 (endogenous control) associated with PAPD2 (Fig. 2E). The result is shown as a fold enrichment of pre-miRNAs over control IgG immunoprecipitates. Pre-miR-125b was enriched over 40-fold in PAPD2 immunoprecipitates (Fig. 2E), while neither pre-miR-133a nor pre-miR-21 (endogenous) was significantly enriched upon pull down of PAPD2 (Fig. 2E). These results demonstrate a specific interaction between PAPD2 and pre-miR-125b, which leads to monoadenylation of miR-125b.

### 3’-end monoadenylaton of miR-125b does not alter stability or subcellular localization

It has been reported that tailing can exhibit an effect on miRNAs’ stability ^13,21,22^ or subcellular localization ^25^. To test whether monoadenylation alters the stability of miR-125b, synthetic RNA oligonucleotides matching the sequence of miR-125b or miR-125b+A were transfected into Cos7 cells, which have a low background of endogenous miR-125b. The amounts of mature miR-125b and miR-125b+A were examined by qRT-PCR 6, 12, 24, and 48 h post-transfection. Throughout the duration of the experiment, there was no difference in the relative expression of either RNA transcript (Fig. 3A), indicating that there is no major difference in miRNA stability between miR-125b and miR-125b+A (Fig. 3A). This result is consistent with the absence of change in the total amount of miR-125b and miR-125b+A upon downregulation of PAPD2 (Fig. 3A).

**Fig. 3.**
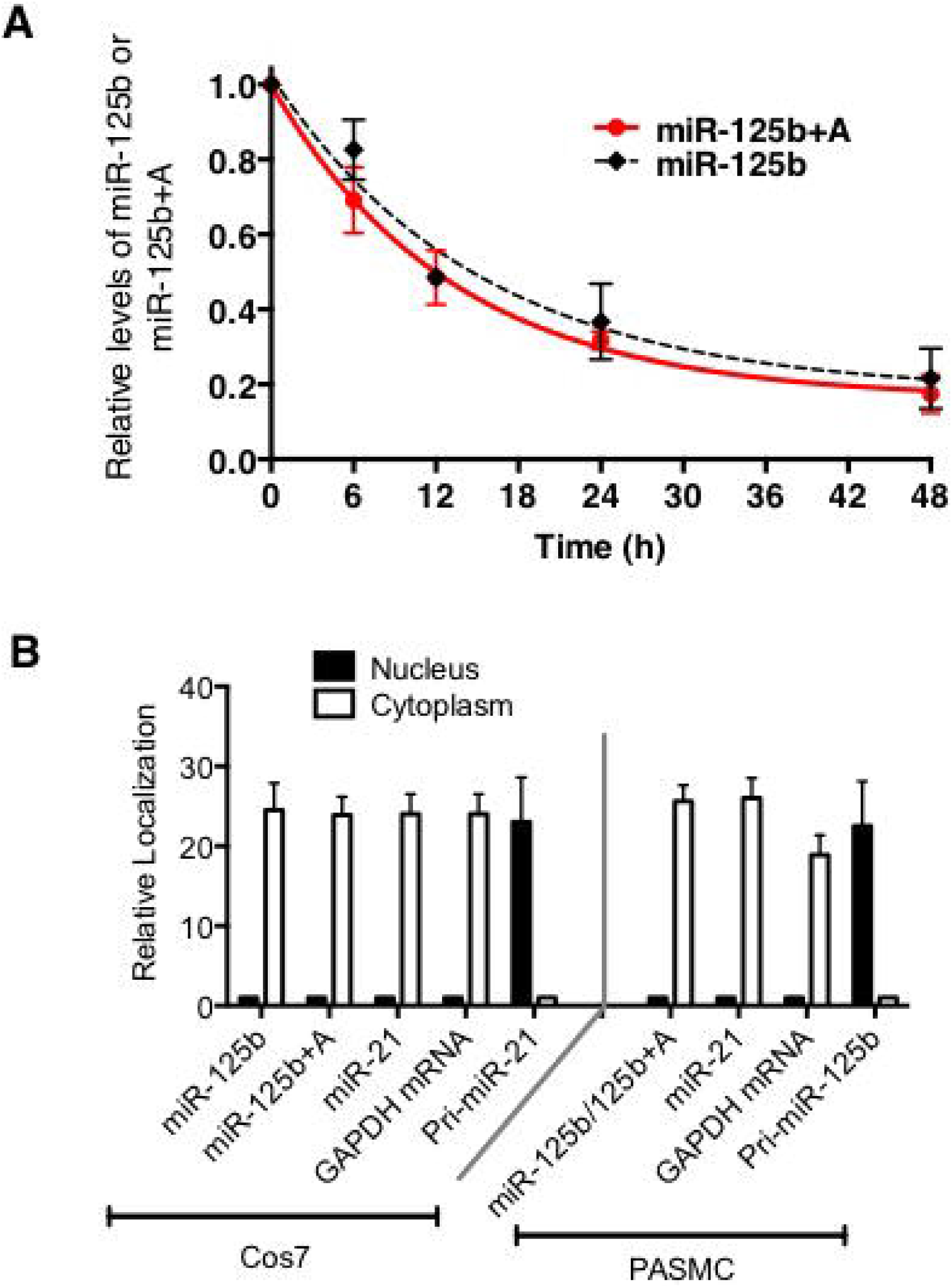
Adenylation of miR-125b does not alter stability or subcellular localization. **A.** Cos7 cells were transfected with unmodified ribonucleic acids with miR-125b or miR-125b+A sequence for 24 h prior to changing media. RNA samples were collected 6, 12, 24, or 48 h after transfection and levels of miR-125b (black dotted line) or miR-125b+A (red solid line) were measured by qRT-PCR analysis. **B.** Cos7 cells were transfected with mock, miR-125b or miR-125b+A for 24 h. Total RNAs were isolated from cytoplasmic or nuclear fraction of Cos7 cells or PASMC, followed by qRT-PCR analysis of miR-125b, miR-21, GAPDH mRNA (control for cytoplasmic RNAs) and pri-miR-21 or -125b (control for nuclear RNAs). RNAs enriched in the cytoplasm (miR-125b, miR-125b+A, miR-21, and GAPDH mRNA) are shown relative to their amount in the nuclear fraction. Conversely, RNAs enriched in the nucleus (pri-miR-125b and pri-miR-21) are normalized to their amount in the cytoplasmic fraction.

Next we examined the subcellular localization of miR-125b by fractionation of the cytoplasm and nucleus of Cos7 cells transfected with miR-125b or miR-125b+A, followed by qRT-PCR analysis. Subcellular fractionation showed that both miR-125b and miR-125b+A are primarily localized in the cytoplasm of Cos7 cells, similarly to miR-21 and GAPDH mRNA (Fig. 3B), unlike pri-miR-21, which is predominantly in the nucleus (Fig. 3B). Similarly, endogenous miR-125b/miR-125b+A (undistinguishable by qRT-PCR) were detected predominantly in the cytoplasm in PASMC, suggesting that both (and in particular miR-125b+A, which is the predominant isoform in PASMC) are localized in the cytoplasm. Since, like the stability, the subcellular localization of miR-125b+A and mir-125b is not significantly different (Fig. 3B), we sought to investigate whether target silencing might differ between miR-125b isoforms.

### Differential mRNA targeting between miR-125b and miR-125b+A

To identify potential targets differentially regulated by *miR-125b and* miR-125b+A, mock, miR-125b or miR-125b+A RNA mimics were transfected into Cos7 cells and the total transcripts were collected 24 h later for next-generation sequencing. Both miR-125b and miR-125b+A mimic exhibit a minimal detectable effect on all cellular transcripts as assessed by the cumulative distribution of the fold-change in transcript expression *vs.* control (mock) treated cells (Fig. 4A, dotted lines). Predicted miR-125b targets by Targetscan showed a significant decrease in mean expression levels, suggesting that both miR-125b mimics silenced a large number of predicted miR-125b targets (Fig. 4A, solid lines). Importantly, there was no difference between the overall degree of silencing by miR-125b and miR-125b+A (*p* >0.05 by ANOVA) – thus both sequences silenced a similar number of predicted targets to a similar degree (Fig. 4A).

**Fig. 4.**
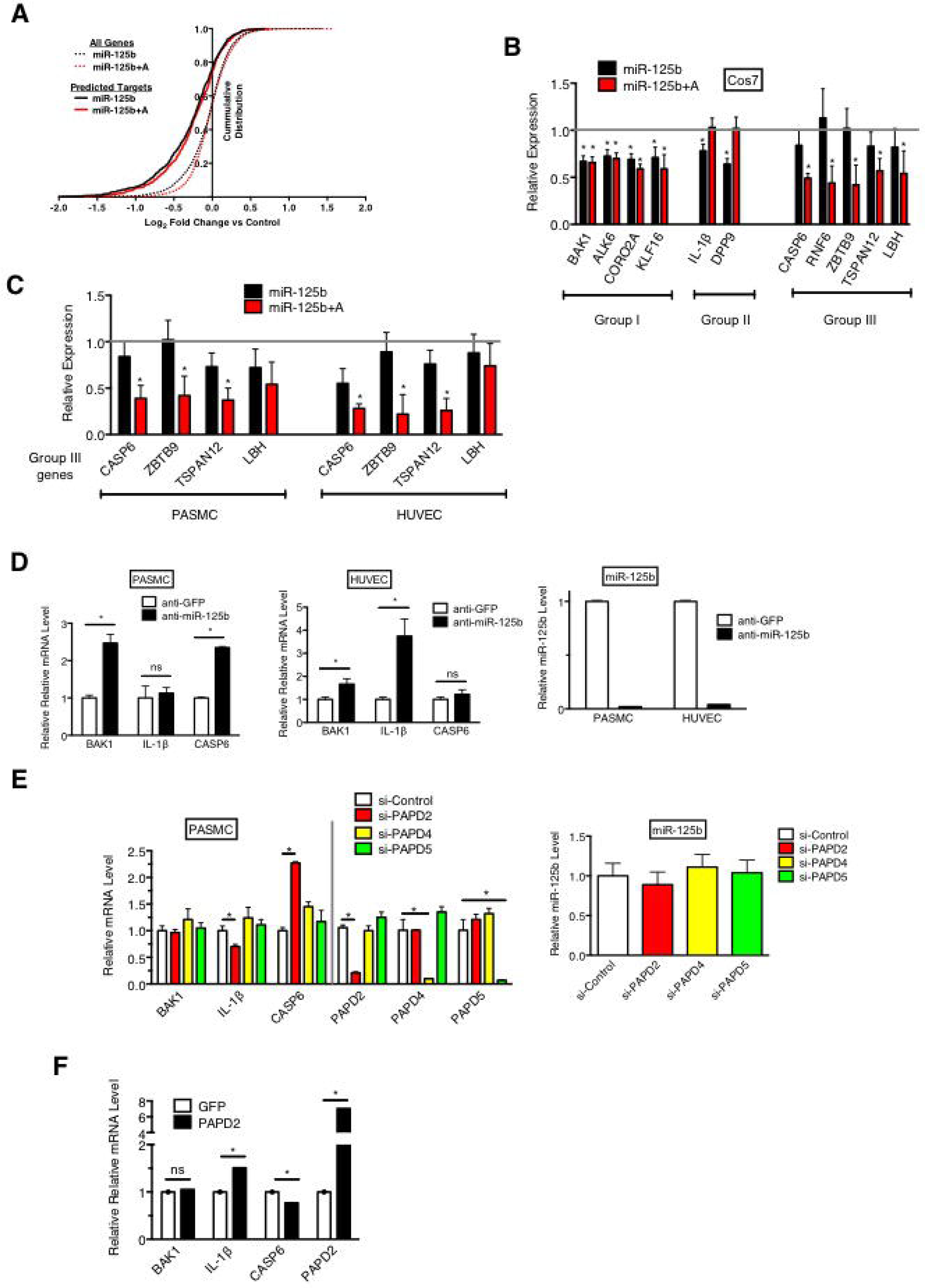
Differential silencing by miR-125b and miR-125b+A. **A.** Cos7 cells were treated with miR-125b or miR-125b+A mimics for 48 h, followed by miR-Seq. The cumulative distribution of gene expression show a significant shift towards inhibition of predicted miR-125b targets (solid lines) as compared to all other genes (dotted lines) and no significant difference in the extend of inhibition between miR-125b (black) and miR-125b+A (red) genes. **B.** Silencing of genes in Group I-III by miR-125b or miR-125b+A are examined in Cos7 cells transfected with miR-125b or miR-125b+A mimic. Bars are mean change in expression (percentage of miR-125b *vs* control) for each gene indicated. Error bars indicate standard error from six independent samples. **C.** Silencing of Group III genes were compared in PASMC and HUVEC transfected with miR-125b or miR-125b+A mimic. Bars are mean change in expression (percentage of miR-125b *vs* control) for each gene indicated. Error bars indicate standard error from six independent samples. **D.** PASMC or HUVEC were transfected with anti-miR-125b or control siRNA (control) for 24 h, followed qRT-PCR analysis of BAK1, IL-1β, and CASP6 mRNAs normalized to GAPDH. Results are shown as mRNA amounts relative to control siRNA-transfected cells (left panel). Efficiency of knock down of miR-125b by anti-miR-125b was examined by qRT-PCR normalized to U6 snRNA (right panel). Results are shown as mean±SEM of three independent experiments. **E.** PASMC were transfected with si-Control, si-PAPD2, si-PAPD4, or si-PAPD5 for 48 h, followed by qRT-PCR analysis of *BAK1*, *IL-1β*, *CASP6*, *PAPD2*, *PAPD4*, and *PAPD5* mRNAs normalized to *GAPDH* (left panel) and miR-125b normalized to U6 snRNA (right panel). Results are shown as mean±SEM of three independent experiments and the relative mRNAs levels are normalized to that of control siRNA-transfected cells (left panel). **F.** PASMC or HUVEC were transfected with anti-miR-125b, anti-miR-143 (control), or anti-miR-100 (control) for 48 h. Caspase-3/7 activity was measured 48 h after transfection. Results are shown as Caspase-3/7 activity relative to mock transfection control and mean±SEM of three independent experiments.

Using a conservative threshold cutoff of a 2-fold decrease in expression as “significantly inhibited”, we detected 125 transcripts inhibited by either or both miR-125b and miR-125b+A mimics (Fig. 4B). We categorized potential targets of miR-125b and/or miR-125b+A into three sets: Group I (equally downregulated by miR-125b and miR-125b+A), Group II (more significantly silenced by miR-125b), and Group III (more significantly silenced by miR-125b+A). Selected members of different Groups of targets were compared by qRT-PCR analysis and generally confirmed the miR-Seq data (Fig. 4C). We thus hypothesized that differentially targeted transcripts might contribute to a PASMC-specific biological effect of miR-125b.

As ∼2/3 of miR-125b in PASMC are monoadenylated, anti-miR-125b treatment in PASMC should derepress both Group I genes (ex. *BAK1*) and Group III genes (ex. *CASP6*), while in HUVEC cells, where miR-125b is mostly unmodified, anti-miR-125b treatment should derepress only Group I genes (ex. *BAK1*) and Group II genes (ex. *IL-1β*). To test this hypothesis, we transfected anti-miR-125b or anti-GFP (control) into PASMC or HUVECs, and measured the amount of *BAK1*, *IL-1β* and *CASP6*. Although anti-miR-125b downregulated the endogenous miR-125b/miR-125b+A similarly in PASMC and HUVEC (Fig. 4D, right panel), its effect on Group II (*IL-1β*) and Group III (*CASP6*) genes was different in PASMC and HUVEC, in contrast with the Group I gene *BAK1*, which was derepressed in both cell types (Fig. 4D). Since the low expression of *IL-1β* mRNA in PASMC at steady state might obscure the effect of anti-miR-125b, we first induced *IL-1β* in PASMC by TNFα treatment, and then assessed the effect of anti-miR-125b *vs* anti-GFP. However, the result was the same as that shown in Fig. 4D and confirmed that induced *IL-1β* was unchanged by anti-miR-125b (Suppl. Fig. S3A).

To provide further evidence that differential regulation of Group II and III genes in PASMC is due to the monoadenylation status of miR-125b, PAPD2 was knocked down by siRNA transfection in PASMC, followed by qRT-PCR analysis of *BAK1* (Group I), *IL-1β* (Group II) and *CASP6* (Group III) (Fig. 4E). As negative controls, we included siRNAs against PAPD4 and PAPD5, which are not involved in the monoadenylation of miR-125b (Fig. 2A and 2B). When PAPD2 was downregulated and the fraction of miR-125b+A reduced to ∼50%, *CASP6* (Group III) was derepressed while *BAK1* (Group I) was unchanged (Fig. 5B), confirming that silencing of Group III genes was less efficient when the miR-125b+A fraction was reduced. Unlike *CASP6*, *IL-1β* (Group II) was repressed further as a result of the increased fraction of canonical miR-125b (Fig. 4E). Conversely, expression of exogenous PAPD2 ∼7-fold over endogenous level led to a derepression of *IL-1β* (Group II) and further decrease of *CASP6* (Group III), while *BAK1* (Group I) was unchanged (Fig. 4F). These results demonstrate that alterations of PAPD2 expression result in a change of the miR-125b:miR-125b+A ratio and modulate target gene expression due to differential sensitivity of Group II and Group III genes to monoadenylated miR-125b.

**Fig. 5.**
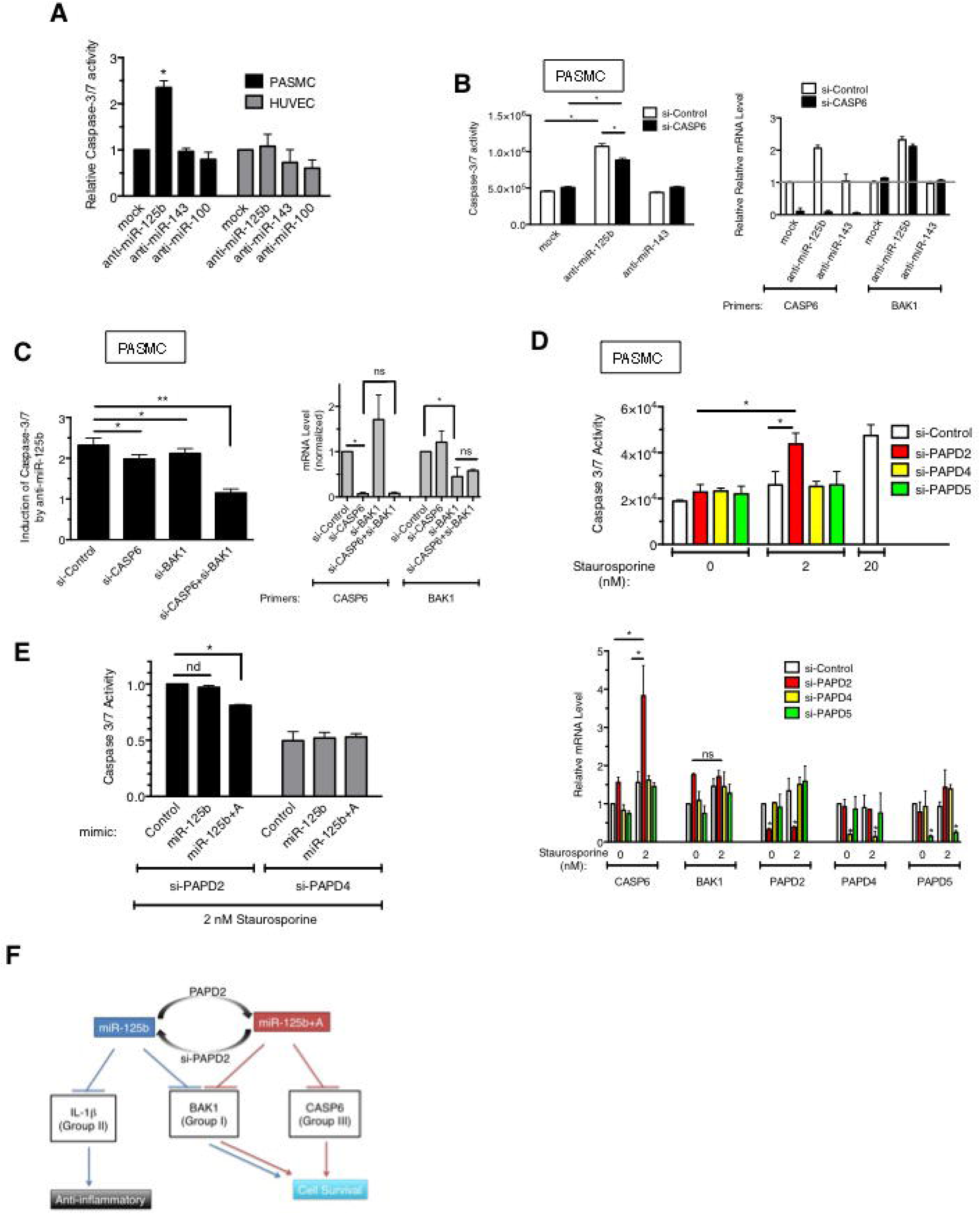
miR-125b isomiR promotes cell survival. **A.** PASMC or HUVEC were transfected with anti-miR-125b or anti-GFP (control) for 24 h, followed qRT-PCR analysis of *BAK1*, *IL-1β*, and *CASP6* mRNAs normalized to *GAPDH*. Results are shown as amount of mRNAs relative to the level of anti-GFP transfected cells (left panel). Efficiency of knock down of miR-125b by anti-miR-125b was examined by qRT-PCR normalized to U6 snRNA (right panel). Results are shown as mean±SEM of three independent experiments. **B.** PASMC were transfected with si-Control (si-GFP) or si-CASP6 for 48 h, followed by Caspase-3/7 assay (left panel). Total RNAs were prepared from the cell and subjected to qRT-PCR analysis of *CASP6* and *BAK1* mRNA normalized to *GAPDH*. Results are shown as mean±SEM of three independent experiments and the relative levels are normalized to that of mock transfected cells (right panel). **C.** PASMC were transfected with si-Control, si-CASP6, si-BAK1, or a combination of si-CASP6 and si-BAK1 in addition to anti-miR-125b or control siRNA (control) for 48 h. Caspase-3/7 activity was measured 48 h after transfection. Results are shown as mean±SEM of fold increase of Caspase-3/7 activity upon anti-miR-125b transfection over control siRNA transfection based on three independent experiments (left panel). Total RNAs were prepared and subjected to qRT-PCR analysis of *CASP6* and *BAK1* mRNA normalized to *GAPDH*. Results are shown as mean±SEM of three independent experiments normalized to si-Control transfected cells (right panel). **D.** PASMC were transfected with si-Control, si-PAPD2, si-PAPD4, or si-PAPD5 for 48 h, followed by treatment with 2 nM staurosporine (ST) for 48 h and the measurement of Caspase-3/7 activity (left panel). As control, si-Control siRNA-transfected cells were also treated with 2 nM ST. Total RNA was extracted and subjected to qRT-PCR analysis of *CASP6*, *BAK1*, *PAPD2*, *PAPD4*, and *PAPD5* mRNA normalized to *GAPDH*. Results are shown as mean±SEM of three independent experiments and the relative levels are normalized to mock-treated, si-Control siRNA-transfected cells (right panel). **E.** PASMC were transfected with si-PAPD2 or si-PAPD4 with control, 15 nM miR-125b, or miR-125b+A mimic for 48 h, followed by treatment with 2 nM ST for 48 h and the measurement of Caspase-3/7 activity. **F.** Schematic diagram of 3’-monoadenylation of miR-125b and its cellular function through differential transcript targeting.

### Derepression of Caspase-6 leads PASMC to apoptotic cell death

To examine the functional significance of miR-125b+A in PASMC, we transfected anti-miR-125b or control anti-miRs (anti-miR-143 and anti-miR-100) into PASMC or HUVEC and measured their effect on cell death by Caspase-3/7 assay. Twenty-four hours after anti-miR-125b transfection, Caspase-3/7 activity began to increase in PASMC (Suppl. Fig. S3B) and the difference between anti-GFP- and anti-miR-125b-transfected cells became significant at 48 h (Fig. 5A, PASMC and Suppl. Fig. S3B). Conversely, anti-miR-125b did not induce apoptosis in HUVEC up to 48 h post-transfection (Fig. 5A, HUVEC). As *CASP6* (Group III) is distinctly downregulated by miR-125b+A in PASMC, while the other pro-apoptotic gene *BAK1* (Group I) is repressed by miR-125b both in HUVEC and PASMC, we hypothesized that anti-miR-125b leads PASMC (but not HUVEC) to apoptosis by removing inhibition of both *CASP6* and *BAK1*. Overexpression of CASP6 was sufficient to induce apoptosis in PASMC (Suppl. Fig. S5). We then examined whether downregulation of CASP6 was able to rescue anti-miR-125b-mediated apoptosis in PASMC (Fig. 5B). PASMC were transfected with anti-miR-125b or anti-miR-143 (control) and si-CASP6 or si-Control (GFP) (Fig. 5B, left panel). The amount of *CASP6* and *BAK1* mRNAs was also measured (Fig. 5B, right panel). As expected, anti-miR-125b, but not anti-miR-143 (control), induced the activation of Caspase-3/7, and the activity was significantly reduced when *CASP6* was targeted by RNAi (Fig. 5B, left panel). However, we observed that BAK1 is activated ∼2-fold by anti-miR-125b and is not downregulated by si-CASP6 (Fig. 5B, right panel), which could potentially reduce the pro-apoptotic effect of si-CASP6. Indeed, when both *CASP6* and *BAK1* are inhibited by RNAi, the induction of Caspase-3/7 by anti-miR-125b is more potently reduced (Fig. 5C). Taken together, these results suggest that monoadenylation of miR-125b by PAPD2, which leads to more efficient silencing of Group III targets, such as *CASP6*, contributes to cell survival in PASMC.

We tested this hypothesis by knocking down PAPD2 in PASMC and measuring their sensitivity to a pro-apoptotic stimulus, such as staurosporine (ST). Typically, 20 nM ST is used to mediate apoptosis in PASMC ^31^. We determined that a concentration of ST reduced to 2 nM was unable to induce apoptosis (Fig. 5D). Under these conditions, we compared the effect of downregulation of different PAPDs on Caspase-3/7 activity (Fig. 5D). Induction of Caspase-3/7 activity was observed only in the cell transfected with si-PAPD2 but not in si-PAPD4 or -5 transfected cell (Fig. 5D, top panel). Furthermore, potent induction of CASP6 (Group III) was observed in si-PAPD2 transfected cells (Fig. 5D, bottom panel), consistent with the result in Fig. 5B. Thus, knock down of PAPD2 sensitizes PASMC to an apoptotic stimulus due to a shift from miR-125b+A to miR-125b and aberrant increase of pro-apoptotic CASP6 (Fig. 5D). Transfection of miR-125b+A mimic rescued the induction of low ST-mediated apoptosis (Fig. 5E), further confirming de-repression of CASP6 as the underlying cause of low ST-mediated apoptosis in si-PAPD2 cells (Fig. 5E). Altogether, our study indicates that a high fraction of miR-125b+A in PASMC provides a protective effect from apoptotic stimuli (Fig. 5F).

### Differential target site selection by miR-125 isoforms

To investigate the mechanism by which adenylation improves Group III genes targeting by miR-125b, we subcloned the full-length 3’ UTR of human *CASP6* (701-nt) into the miRNA targeting luciferase reporter pIS0^32^ to generate pIS0-FL (Fig. 6A, “FL”). *CASP6* had been previously identified as a confirmed target of miR-125b, but the targeted site had not been reported^33^. According to a miRNA target prediction algorithm [miRanda, as reported by miRTarBase^34^ (http://mirtarbase.mbc.nctu.edu.tw/php/detail.php?mirtid=MIRT002375#target)], three potential miR-125b recognition elements (MRE1-3) were predicted in the *CASP6* 3’ UTR (Fig. 6A, sequences aligned to miR-125b. Blue characters are conserved among vertebrates). Thus, we generated constructs with three-nucleotide mutations in the region corresponding to the seed sequence of each site (Fig 6A, Mut1-Mut3). Co-transfection of these luciferase reporters and miR-125b, miR-125b+A or miR-100 (control) mimics in Cos7 cells, followed by luciferase assay, was used to measure targeting of *CASP6* UTR by miR-125 isoforms (Fig. 6B). As observed with endogenous *CASP6* mRNA in PASMC and Cos7 cells, the FL construct recapitulated a preferential targeting by miR-125b+A compared to miR-125b (Fig. 6B). *CASP6* UTRs with mutations in MRE1 (FL-Mut1) and MRE3 (FL-Mut3) retained this differential sensitivity to miR-125 isoforms, but the FL-Mut2 could not be inhibited by either miR-125b or miR-125b+A (Fig. 6B), suggesting that the MRE2 (nt 502-519 of *CASP6* 3’ UTR) is necessary for miR-125b/miR-125b+A-dependent repression (Fig. 6B, FL-Mut2). To investigate whether MRE2 is sufficient for downregulation by miR-125b/miR-125b+A, we cloned the sequence containing MRE1 (S1, 30-nt), MRE2 (S2, 26-nt), or MRE3 (S3, 30-nt) individually into the pIS0 vector and performed a luciferase assay (Fig. 6C). While S1 and S3 were not inhibited by either miR-125b isoform, S2 was targeted equally by miR-125b and miR-125b+A (Fig. 6C). Thus, we confirmed that MRE2 is a *bona fide* miR-125b recognition site, but MRE2 alone did not produce the incremental inhibitory effect seen with miR-125b+A on the full-length *CASP6* 3’ UTR. To investigate the discrepancy between FL vs S2, we hypothesized that miR-125b+A and miR-125b might target overlapping but distinct sequences in the *CASP6* 3’ UTR. In order to discover potential differences in miRNA-*CASP6* 3’ UTR structures between miR-125b isoforms, we employed a bimolecular folding algorithm [“bifold”, a component of the RNAstructure Web Server^35^, (http://rna.urmc.rochester.edu/RNAstructureWeb/Servers/bifold/bifold.html)], which predicts the lowest free energy structures for two interacting sequences, allowing intramolecular base pairs. A 55-nt long extended version of S2 (which we name LS2-WT, corresponding to nucleotides 455-519 of the *CASP6* 3’ UTR sequence) could be folded with miR-125b into a bimolecular structure analogous to the match predicted by miRanda for S2 (Fig. 6D*a*, compare to the aligned nucleotides in S2, Fig. 6A). The free energy released by this predicted folded structure is E=-17.1. However, several more thermodynamically favored structures can be predicted for the same pair of RNAs. Among those that preserve the seed pairing for S2, the lowest free energy fold (E=-18.4) for miR-125b+A and LS2-WT is shown in Fig. 6D*b*. We notice that this structure allows an additional A-U base pair for the 3’ adenine in miR-125b+A (Fig. 6D*b*, red circle), further stabilizing this structure compared to folding with canonical miR-125b. Both structures would be disrupted by the M2 mutation, as they share the same seed sequence (Fig. 6D, “Seed match”), but they use different “3’ supplementary sites” (Fig. 6D, “Suppl. pairing”)^36^. To test whether this alternative conformation may contribute to miR-125b+A targeting of *CASP6* 3’ UTR, we generated two new luciferase reporter constructs. One contains the correct LS2-WT sequence downstream of the luciferase gene; the other has the same region with 10 substitutions in the alternative 3’ supplementary site (Fig. 6D, bottom panel, LS2-Mut). In a luciferase assay performed after transfection of these constructs into Cos7 cells, LS2-WT recapitulated the behavior of FL in response to miR-125b isoforms, with significant inhibition by canonical miR-125b, and further repression by miR-125b+A (Fig. 6E, LS2-WT). Conversely, although the LS2-Mut and S2 constructs were still inhibited by miR-125b, they were not further repressed by miR-125b+A (Fig. 6E, LS2-Mut). We interpret this result as an indication that the alternative 3’ supplementary site (Fig. 6D) plays an important role in the targeting of *CASP6* 3’ UTR by miR-125b+A.

**Fig. 6.**
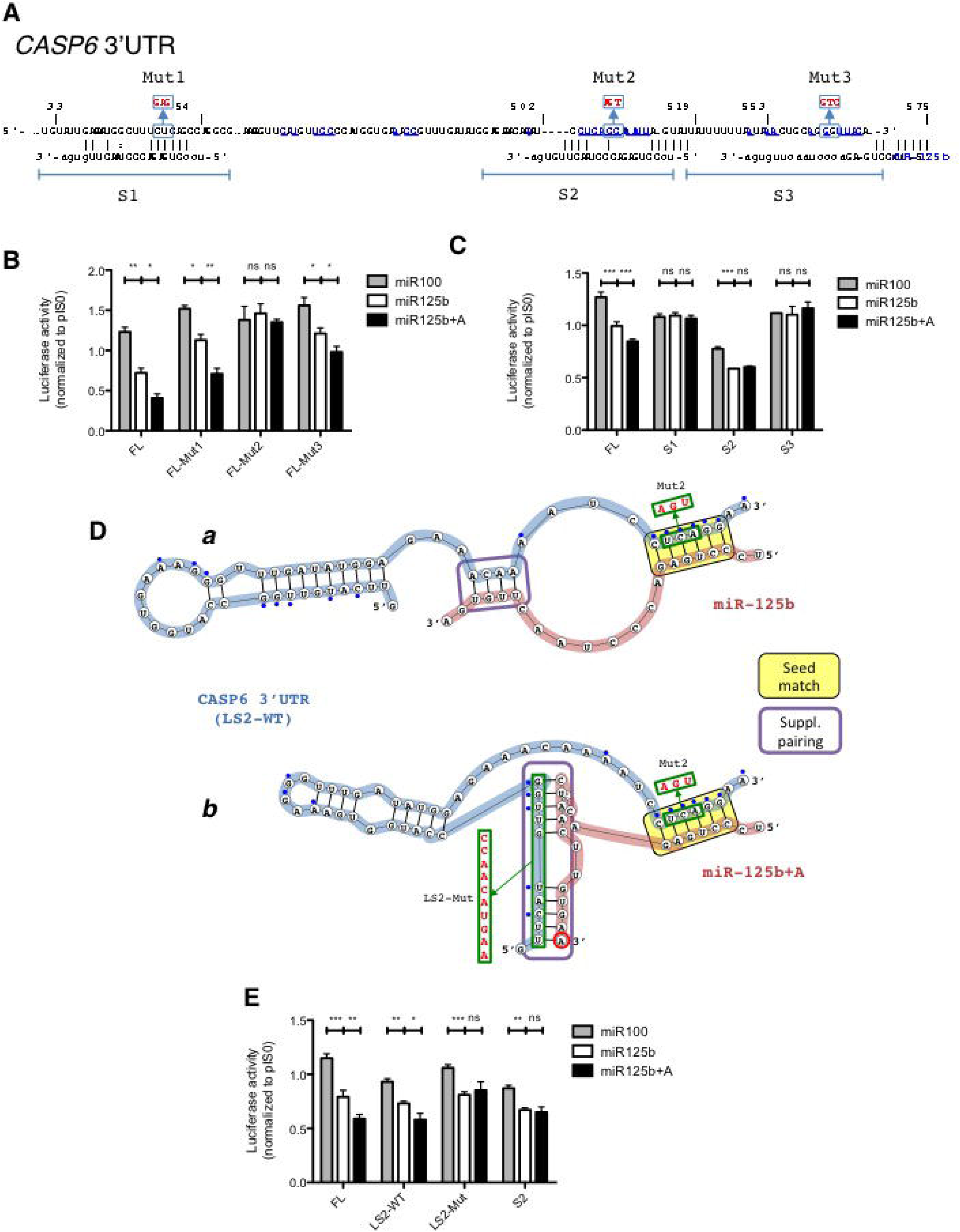
Mechanism of improved silencing of *CASP6* by miR125b+A. **A.** Scheme of the *CASP6* 3’ UTR. The partial sequence shown (numbered from the beginning of the 3’ UTR) includes the MRE1-3 for miR-125b as predicted by the miRanda algorithm. Three-nucleotides mutations (Mut1-3) are indicated by blue boxes. Sites containing the MREs are labeled S1-3. Nucleotides underlined and labeled in blue are conserved among vertebrates. **B.** Cos7 cells were transfected with luciferase reporter plasmids derived from pIS0, as described, including the entire *CASP6* 3’ UTR (FL) and the Mut1-3 mutant constructs, together with miR-125b, miR-125b+A or miR-100 (control) mimics for 48 h. A luciferase assay was used to measure constructs expression. Results are shown as mean±SEM of three independent experiments and are normalized to empty pIS0 vector-transfected cells. **C.** As in B., comparing the entire *CASP6* 3’ UTR (FL) with single MRE constructs S1-3. **D.** Scheme of the predicted structure of the LS2-WT sequence from *CASP6* 3’ UTR (from nucleotide 455 to 519, blue ribbon) in conjunction with miR-125b (***a***) or miR-125b+A (***b***) (red ribbons), as folded by the “bifold” algorithm. Annotations include: Mut2 and LS2-Mut sequences (green boxes), seed match sequence (yellow background box), 3’ supplemental pairing sequence (purple-lined box), miR-125b+A adenylation site (red circle). **E.** As in B., comparing the entire *CASP6* 3’ UTR (FL) with constructs LS2-WT, LS2-Mut and S2.

## Discussion

In this study, we report a high rate of 3’-monoadenylation of one of the most abundantly expressed miRNAs in human and rat PASMC. No other tissue expresses miR-125b+A to a level comparable to PASMC, suggesting that miRNA monoadenylation can play an important tissue-specific role. Genome-wide analysis of miR-125b and/or miR-125b+A target mRNAs revealed a set of miR-125b targets that are more efficiently silenced by miR-125b+A than miR-125b. We found that one such target, CASP6, is a potent inducer of apoptosis in PASMC. Conversion of miR-125b+A to miR-125b in PASMC derepresses CASP6 expression and sensitizes PASMC to an apoptotic stimulus. Thus, we demonstrate that the 3’-end monoadenylation of miR-125b is a novel mechanism of protection of PASMC from cell death.

In addition to 3’-adenylation, mature miRNAs and pre-miRNAs undergo 3’-mono- or oligo-uridylation, which affects miRNA stability and activity. Both ZCCHC11 (also known as TUT6) and ZCCHC6 (also known as TUT6) uridylate pre-let-7 through interaction with Lin 28 and control pre-let-7 stability and processing to mature let-7 ^14,16,18^. ZCCHC11-mediated uridylation of miR-26 modulates its activity ^20^. Remakably, proteins that localize primarily to the nucleus ^37^, such as ZCCHC11^20^, PAPD2 ^38^, PAPD5 ^13^ and PAPD4 ^21,22^, can modify miRNAs that are in the cytoplasm. We show that the 3’-monoadenylation reaction does not occur when mature miR-125b is introduced in cells, but ∼35% of miR-125b is monoadenylated when pri-miR-125b is expressed. Taken together with the specific interaction between PAPD2 and pre-miR-125b and the predominant expression of PAPD2 in the nucleus, we propose that PAPD2 is loaded on pri- or pre-miR-125b in the nucleus prior to translocation to the cytoplasm and Dicer processing, whereupon it catalyzes the 3’-adenylation of miR-125b.

PAPD2 was cloned and originally named Terminal Uridylyl Transferase 1 (TUT1) based on its function as human U6 snRNA-specific terminal uridylyl transferase ^38^. Later, Laishram and Anderson discovered a poly(A) polymerase (PAP) activity in TUT1, which is required for 3’-end cleavage and polyadenylation of a select set of pre-mRNAs (mRNAs), including heme oxygenase (HO-1) mRNA, and renamed the protein Star-PAP^39^. Nontemplated 3’-monoadenylation of miRNAs by PAPD2 has not been reported previously. However, downregulation of PAPD2 by siRNA in human colon carcinoma HCT-116 cells results in a global decrease of miRNA levels by approximately 40%, as measured by qRT-PCR-based miRNA profiling ^40^. The mechanism of this global miRNA suppression appears to be independent of the nucleotidyl transferase activity of PAPD2 as it occurred irrespectively of changes in 3’-tailing ^40^. Our miR-Seq analysis using RNAs from PASMC transfected with siRNA against PAPD2, PAPD4, or PAPD5 indicates that downregulation of PAPD2 does not mediate global downregulation of miRNAs in PASMC because total miRNA read counts do not significantly differ from those from si-Control, si-PAPD4, or si-PAPD5-transfected PASMC (Suppl. Table 2). The discrepancy between the previous report and our study might be due to the difference in tissue type (colon vs vascular smooth muscle), malignant vs non-malignant cells, or miRNA quantitation methodology (qPCR vs miR-Seq).

Several PAPDs participate in nontemplated 3’-tailing of miRNAs. One example is the 3’-monoadenylation of miR-122 by PAPD4 (also known as GLD2). miR-122 is a liver-specific, highly abundant miRNA, and PAPD4-mediated 3’-monoadenylation increases its stability ^21,22^. PAPD5 catalyzes the 3’-monoadenylation of a miR-21+C isomiR (23-nt, containing a templated C at the 3’-end of the canonical 22-nt miR-21 sequence), to produce the miR-21+CA isomiR (24-nt), more stable than its unmodified counterpart in human cancer cells MCF7 and THP1 ^13^. As this study reports only the adenylation ratio (miR-21+CA reads count/miR-21+C reads count), the abundance of miR-21+C or miR-21+CA in comparison with canonical miR-21 is unclear ^13^. In PASMC, we found that ∼66% and 2.2% of total miR-21 reads correspond to miR-21+C and miR-21+CA, respectively (data not shown), pinning the adenylation ratio to 0.07, significantly lower than that of miR-125b (∼2.5). Furthermore, our results indicate that PAPD5 knockdown in PASMC or overexpression of PAPD5 in human breast carcinoma MCF7 do not alter the abundance of miR-125b+A ^13^ (see Suppl. Fig. S4). Thus, we propose that the 3’-monoadenylation of miR-125b is specifically catalyzed by PAPD2 and not by PAPD5, underscoring the miRNA-specific activity of PAPD2 and PAPD5.

The mechanism by which miRNAs target their cognate mRNAs has been the focus of intense study. A conspicuous number of miRNA/mRNA interactions can be explained by the “seed match rule”, by which a short consecutive match toward the 5’ end of the miRNA initiates a miRNA-target duplex, and subsequently partial annealing may propagate to further stabilize the miRNA-target hybridization ^36^. The *CASP6* 3’ UTR site recognized by miR125b (MRE2) contains a 6-mer match to position 3–8 that has been classified as an “offset 6-mer seed” because of its position and its often marginal effect on repression ^41^. However, transcriptome-wide miRNA binding studies have determined that perfectly matched miRNA seeds are neither necessary nor sufficient for all functional miRNA-target interactions, and indeed about 60% of miRNA-binding activity does not follow the canonical rule about the seed matching between miRNA and target mRNAs ^42,43^. When a biologically validated target site contains a seed sequence that is imperfect or suboptimal, a “3’ supplementary (or compensatory) site” or “seed-like motif” is often found to counterweigh the deficiency in the miRNA-target hybridization stability ^44^. Based on the occurrence in predicted and validated sites, it has been proposed that the distance between seed match and 3’ supplementary site vary between 1 and 5 nt ^36^. However, no experimental evidence indicates that this is the maximal size of the loop. In the *CASP6* 3’ UTR, we detect a role for a 3’ supplementary site located 36 nt away from the offset 6-mer seed sequence. More data on other non-canonical miRNA targeting is required to gauge how common this feature is to other miRNA-mRNA structures. Additionally, imperfect miRNA targeting has been found in situations in which miRNA expression is spatially or temporally regulated, as in the case of let-7 and lin-41 ^36^. It is conceivable that a site able to sense the presence of a single additional A at the 3’ end of a miRNA might be highly atypical and poised to switch between alternative structures. The miR-125b site in *CASP6* and other Class III genes might have been selected to precisely respond to the tissue-specific adenylation status of miR-125b, and therefore provide a prototype for a new type of miRNA sites.

Indeed, it is known that miR-125b exhibits several context-dependent activities ^30^. For example, miR-125b can act as both a tumor suppressor and an oncogene ^30^. Furthermore, aberrant expression of miR-125b is associated with diverse pathologies, including malignancies ^45-48^ and vascular inflammation and calcification^28,29^. Our study suggests that the context-dependent activities of miR-125b can be, in part, due to a spatial and temporal change in the fraction of miR-125b+A. Furthermore, a change in the miR-125b+A fraction could occur under pathological conditions and contribute to the development of disorders. We demonstrate that the miR-125b+A/miR-125 ratio is essential for PASMC survival. When overall miR-125b+A and miR-125b levels are reduced, PASMC undergo apoptotic cell death that can be partially rescued by downregulating CASP6 by siRNA; this suggests that elevation of CASP6 level contributes to the activation of the apoptotic cascade. Indeed, exogenous expression of CASP6 induces apoptosis in PASMC. CASP6 is a member of the Cysteine-aspartic acid protease (Caspase) family. CASP6, along with Caspase-3 (CASP3) and Caspase-7 (CASP7), is an apoptotic effector activated by CASP3 ^49^ and thought to function as a downstream enzyme in the caspase activation cascade ^49^. CASP6 also undergoes self-processing without other members of the caspase family ^49^. Aberrant expression of *CASP6* and mutations in *CASP6* are associated with different types of cancer ^50-53^, while anomalous activation of CASP6 is linked to neurodegenerative diseases, such as Huntington’s and Alzheimer’s disease ^49^. Generally, malignant cells express a higher fraction of monoadenylated miR-21+C compared to normal cells ^13^. Similarly, aberrant expression and/or activity of CASP6 in neurodegenerative conditions might be due to a change in the fraction of miR-125b+A isomiR. Finally, our study sheds lights on a potential link between aberrant expression or activity of CASP6 and vSMC proliferative conditions, such as pulmonary artery hypertension and restenosis. It is plausible that a change in miR-125b+A/miR-125b ratio could contribute to the pathogenesis of those vascular diseases. Although we developed a new LD assay to detect miR-125b+A isomiR semi-quantitatively, this assay is not readily applicable to 3’-tailings of other miRNAs and its sensitivity and quantitative power are limited. Future development of a rapid, sensitive, and accurate isomiR quantitation method will allow measuring the isomiR level in primary tissue samples and addressing its significance in pathogenesis.

## Methods

### Cell Culture and growth factor stimulation

Human primary pulmonary arterial smooth muscle cells (PASMC) were obtained from Lonza (CC-2581) and maintained for no more than 8 passages in Smooth Muscle Cell Growth Media (SM-GM2, Lonza) supplemented with 5% FBS and 1X Penicillin/Streptomycin (Gibco). Endothelial cells were maintained in Endothelial Cell Basal Media (Lonza) supplemented with the complete Endothelial cell singlequot nutrients (Lonza). H1299 cells were maintained in RPMI containing 10% FBS (Hyclone) and 1X Pen/Strep. All other cells were cultured in DMEM supplemented with 10% FBS and 1X Pen/Strep. Prior to growth factor stimulation, cells were serum starved for overnight in DMEM containing 0.2% FBS, followed by treatment with 3 nM BMP4(R&D Systems), 50 pg/ml TGF-β1 (R&D Biosystems), or 20 ng/ml PDGF-BB (R&D Systems), TNFα stimulations (10 ng/ml, R&D Systems) or staurosporine (2 or 20 nM, Sigma).

### siRNA and plasmid transfection

Unless otherwise indicated, siRNAs were synthesized by Dharmacon and transfected into cells at a concentration of 100 nM in DMEM/0.2% FBS media using RNAiMax (Life Technologies). Cells were treated with siRNA for 24 h prior to growth factor treatment and additional 24-40 h of culture. The following siRNA sequences were used to knockdown the indicated genes: siPAPD1 5’ - GCUGGUAUCCCUAUUGCUA - 3’, siPAPD2 5’ - UGACCUUGCUGGUGAUCUA - 3’, siPAPD4 5’ - GAGCAGUGAUGGUGAUUUA - 3’, siPAPD5 5’ - GAAUAGACCUGAGCCUUCA - 3’, siPAPD7 5’ - CAGACAAGCUGGUGUAGAA - 3’, siGFP 5’ - AAGGCAAGCUGACCCUGAAGU - 3’ (2’-O methylated for use as control in miRNA mimic experiments). siGenome Control 1 (Dharmacon) was used as a negative control for all siRNA treatment experiments. 2’-O-methylated miRNA mimics matching the exact sequence of miR-125b and the adenylated miR-125b were obtained from Applied Biosystems and transfected into cells at a concentration of 20 nM. Unmodified miRNA oligoribonucleotides were synthesized by IDT and used for the miRNA stability assays as well as for the RNA-Seq experiments. For all mimic experiments siGFP was used as a negative control. Anti-mir-125b was obtained from Applied Biosystems. mPAPD2 was PCR amplified from PASMC cDNA using the following primers: 5’ - GGGGTACCCCGCCACCATGGCGGCGGTGGATTC-3’ and 5’ - GGGAATTCCTCACTTATCGTCGTCATCCTTGTAATCCTTGAGGAGATTTTT-3’. The resulting PCR product was ligated into the KpnI and EcoRI sites of a pcDNA3.1+ vector and their identity was confirmed by sequencing. Cells were transfected with 1 μg DNA per million cells using 3 μg Fugene6 per μg DNA for 24 h.

### miR-Seq, RNA-Seq and Data analysis

Small RNAs were isolated, purified, and subjected to create small RNA libraries using an Illumina Sequencing prep kit. Small RNA libraries were sequenced using HiSeq 2500 (Illumina) at the UCSF Center for Advanced Technology and Beijing Genome Institute (Shenzhen, China). The resulting library was aligned against mirbase v19. Highly expressed miRNA sequences were manually analyzed for non-genomic additions and only sequences aligning perfectly to the published sequence at all except the last base pair were assessed for analysis and used for calculating extent of modifications.

### Ligation and Digestion (LD) Assay

Small RNAs were purified according to manufacturer’s protocol using Trizol (Life Technologies). RNAs ligated to an adenylated 3’-adaptor (5’-TTCTGGCTCGTACCTTATATCTTCCG-3’) using deletion mutant T4 RNA Ligase 2 (Epicenter). Adaptor ligated RNA was then gel purified using a reducing 10% TBE-PAGE Urea-gel. The resulting RNA was reverse transcribed using Super Script 2 Reverse Transcriptase (Life Technologies) per manufacturer’s protocols and the following adaptor-specific reverse primer: 5’-CGGAAGATATAAGGTACGAGCCAGAA-3’. The cDNA was PCR amplified for 25-35 cycles using the AccuPower PCR Mix system (Bioneer) and primers specific to the miRNA of interest – miR-125b 5’-TCCCTGAGACCCTAAC-3’, miR-21 5’-AGCTTATCAGACTGATGTT-3’. The PCR product was then digested for 1 hour at 37°C with EcoRI. The digested DNA was then run on a non-reducing 15% TBE-PAGE gel and visualized using SyberGold.

### Quantitative RT-PCR analysis

Total RNAs were purified using Trizol reagent (Life Technologies) according to manufacturer’s recommendations. 10-50 ng/µl total RNA was first transcribed into cDNA using the first strand cDNA synthesis kit (Thermo Scientific), then quantified by qPCR using the SYBR Green Master Mix (Applied Biosystems). Sequences for all RT-PCR primers (IDT) are shown in Table 1. miRNAs were measured using the TaqMan Array Kits (Applied Biosystems). PCR primers are listed in the file of Supplementary Information.

**Table 1.** Sequence reads number of PASMC miR-Seq analysis. PASMCs were untreated or treated with 3 nM BMP4 for 24 h, followed by small RNA preparation and sequencing library construction. miR-Seq was performed using three independent RNA samples.

| Library | Stimulation | FL | +A | +U | +C | +G | trimmed | Total No.<br>of miR-125b<br>Reads |
| --- | --- | --- | --- | --- | --- | --- | --- | --- |
| **PASC Lib#1** | Untreated | 144296 | 212134 | 9625 | 5681 | 0 | 56966 | 371736 |
|  | +BMP4 | 117395 | 209599 | 7801 | 2863 | 0 | 30252 | 337658 |
| **PASC Lib#2** | Untreated | 1781 | 5887 | 65 | 110 | 0 | 698 | 7843 |
|  | +BMP4 | 1229 | 3647 | 0 | 26 | 0 | 406 | 4902 |
| **PASC Lib#3** | Untreated | 12742 | 72594 | 630 | 1096 | 0 | 5394 | 87062 |
|  | +BMP4 | 7488 | 65738 | 553 | 574 | 0 | 3518 | 74353 |

**Table 2.**
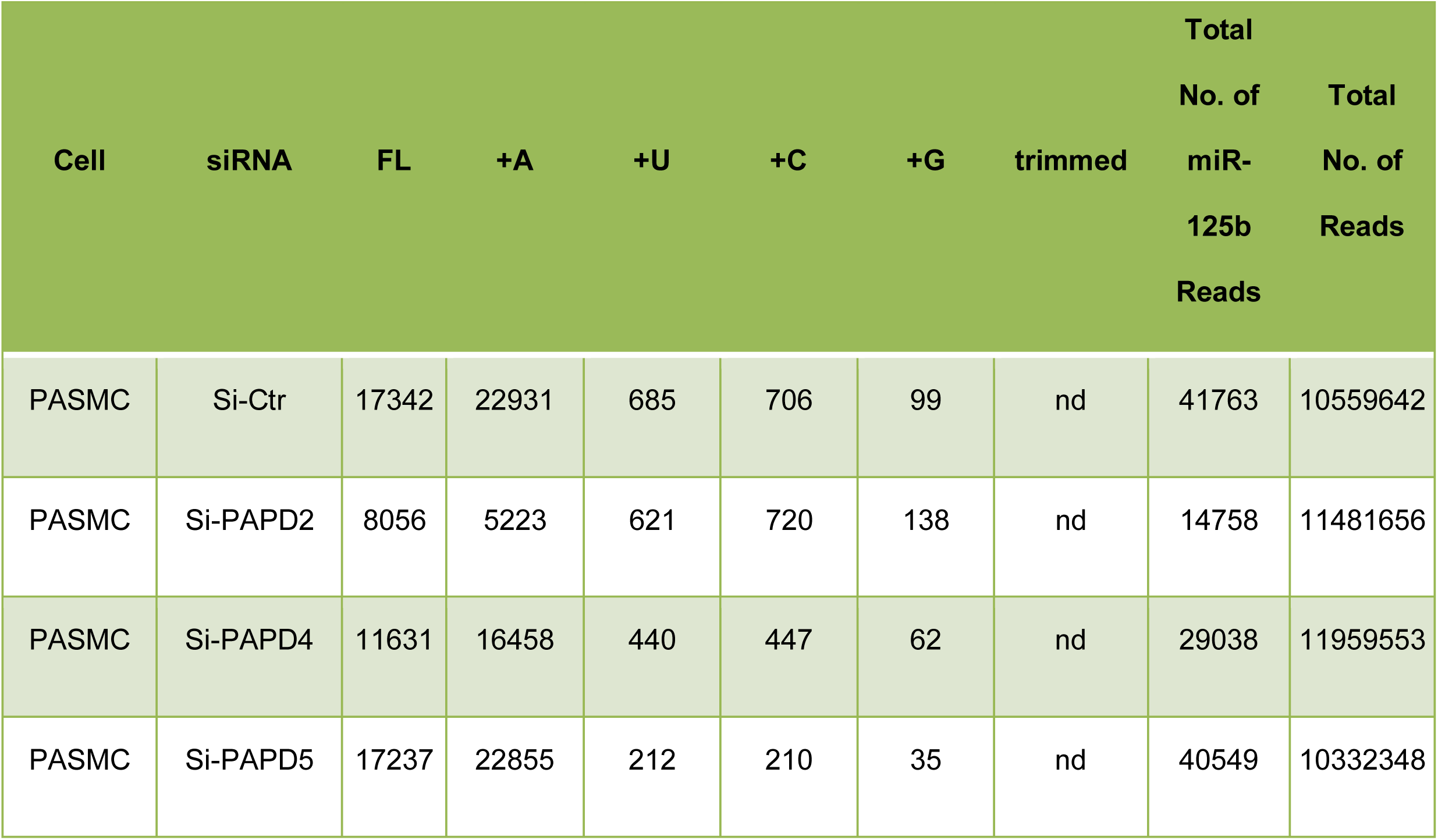
Sequence reads number of PASMC miR-Seq analysis. PASMCs were transfected with si-Control (si-Ctr), si-PAPD2, si-PAPD4, or si-PAPD5 for 24 h, followed by RNA preparation and small RNA library construction.

### Immunoblotting

Cells were lysed by RIPA, and protein concentration was determined by BCA protein assay kit (Sigma, USA). Equal amounts of the extracts were subjected to 8% SDS-PAGE, transferred onto nitrocellulose membranes, and blotted with primary antibodies for PAPD2 (Abcam, Cambridge, MA, USA), smooth muscle actin (SMA, Sigma), anti-Flag (Sigma), anti-GFP (Sigma), and GAPDH (Santa Cruz Biotechnology, Santa Cruz, CA, USA). Immunoblots were subsequently incubated with IR-labeled secondary antibodies (Santa Cruz) and visualized using Image Studio 2.1 (LI-COR USA).

### Luciferase Assay

For creation of miRNA reporter assays, full-length UTRs were PCR amplified or miRNA sites were created by annealing oligos matching the 3’ UTR sequence of interest exactly, and inserted between the SacI and the XbaI sites downstream of the luciferase reporter in a PIS0 construct. Utilized oligo sequences are available upon request. All of the constructs were verified by DNA sequencing.

For luciferase assays, Cos7 cells were seeded in a 6-well plate (2.0×10^5^ per well), co-transfected with 500 ng *CASP6* 3’ UTR and derived firefly luciferase reporter constructs, 5 ng pCMV-Renilla-luciferase reporter vector, and 10 nM miR-125b, miR-125b+A or control RNA 3’O-methyl-modified mimic using 5 μl Lipofectamine 2000 (Life Technologies) in a 500 μl reaction mixture according to the manufacturer’s instructions. Luciferase activity was measured 48 h later in triplicate using a dual luciferase reporter assay system (Promega).

### RNA immunoprecipitation (RIP)

Cos7 cells were infected with adenovirus expressing rat pri-miR-125b or pri-miR-133a, obtained from Dr. E. Olson’s lab ^54^. PASMC were transfected with Flag-tagged human PAPD2 expression vector or GFP expression vector (control) as described above and incubated for another 24 hr. Cross-linking and RNA immunoprecipitation were performed following a published protocol ^55^. Briefly, cells on ice were subjected to four rounds of UV cross-linking using 400 KJ/cm^2^ of 452 nm UV light, then lysed using SDS Buffer (15 mM Tris-HCl, 5 mM EDTA, 2.5 mM EGTA and 1% SDS) and rotating for 30 min at 4 °C. Cell membrane and DNA were disrupted by sonication and the supernatants collected. Immunoprecipitation was performed using anti-flag, M2 conjugated beads (Sigma) overnight at 4 °C. Beads were then sequentially washed with high salt buffer and treated with 0.8 mg/ml Proteinase K for 20 min at 37 °C as described previously^56^. The reaction was stopped with 7 M Urea and centrifuged. The resulting supernatant was collected and treated with DNAse I (Roche) for 20 min at 37 °C. One-tenth of the resulting volume was loaded into each reaction and used for reverse-transcription and qPCR as described above. For input, 1% of original buffer volume was saved and treated with proteinase K and DNAse I, as described.

### Caspase-3/7 Assay

Supernatant medium from treated PASM cells was mixed 1:1 with 25 µL Caspase-Glo 3/7 substrate in a solid white polystyrene 96-well Assay Plate (Corning) and set at room temperature for 30 min prior to reading with a GloMax luminometer (Promega). Integration time was set at 1 second ^54,55^.

### Data availability

The data that support the findings of this study are available within the paper and its supplementary information file, or from the corresponding author on request.

## Acknowledgement

We thank Drs. S. Gnerre, R. Deo, and M. Ku for assisting with the miR-Seq data analysis. We also thank Dr. D. Hart and all members of the Hata and Lagna laboratories for critical discussion. This work was supported by grants from the National Institute of Health: HL093154 and HL108317, and Fondation LeDucq to A.H. and 5T32HL007731-20 to M.T.B.

## Author information

Matthew T. Blahna and Jean-Charles Neel

These authors contributed equally to this work

## Contributions

M.T.B., J.-C.N., G.Y. and G.L. designed and performed experiments and analyzed the results. J.L. and P.G. performed experiments. G.L. and A.H. analyzed and discussed the results, and wrote the manuscript.

