## Supplementary figures and images for "Tissue-specific 3 prime-end adenylation of miR-125b mediates cell survival"

### Supplemental Fig. S1

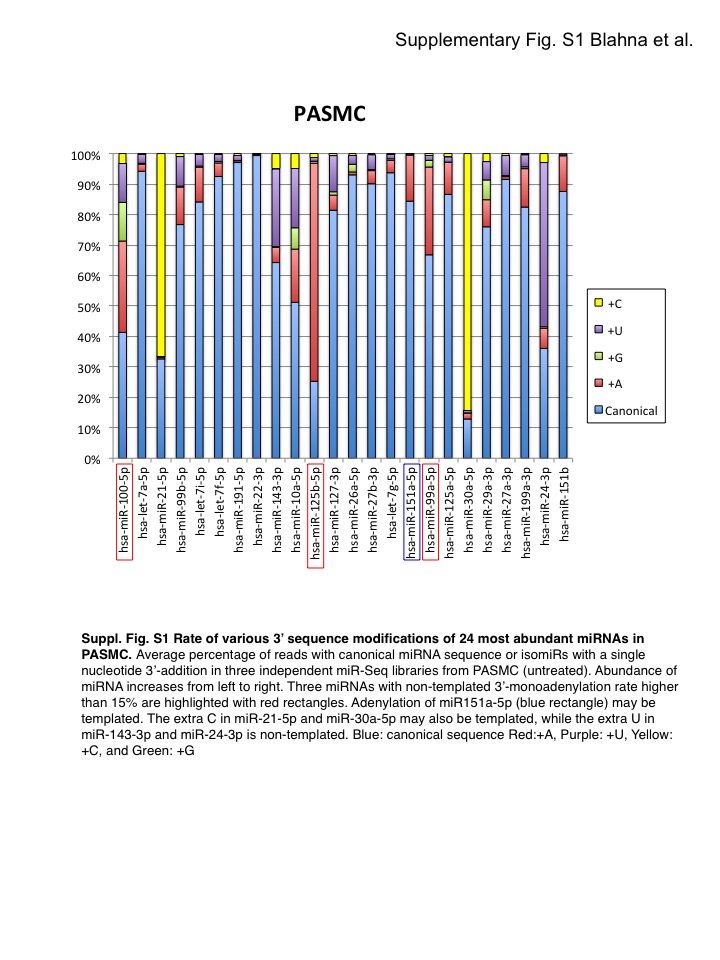

### Supplemental Fig. S2

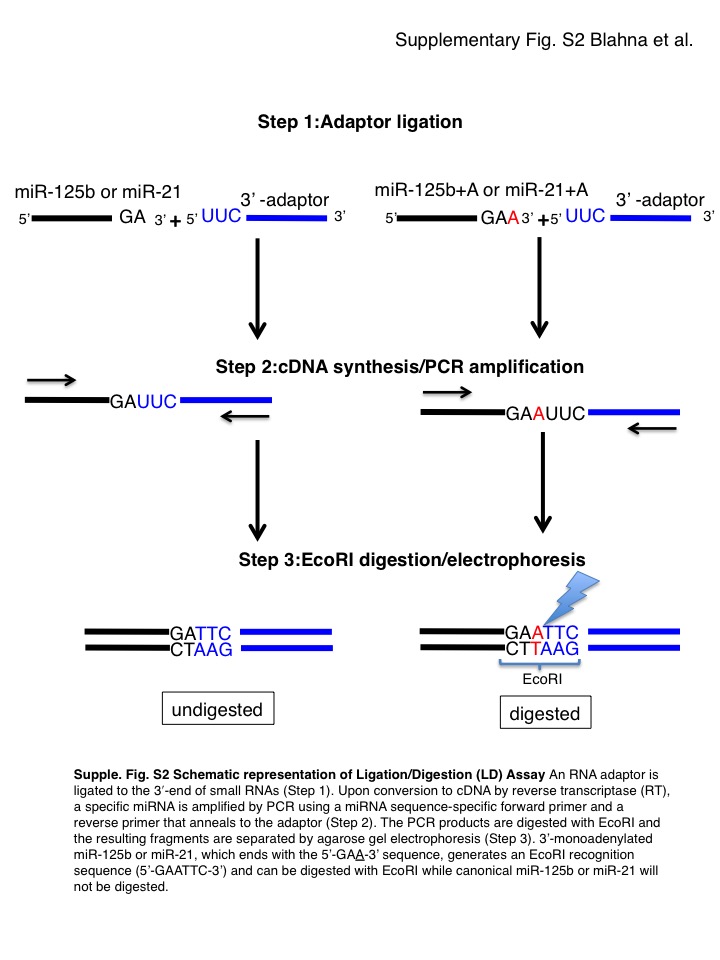

### Supplemental Fig. S3

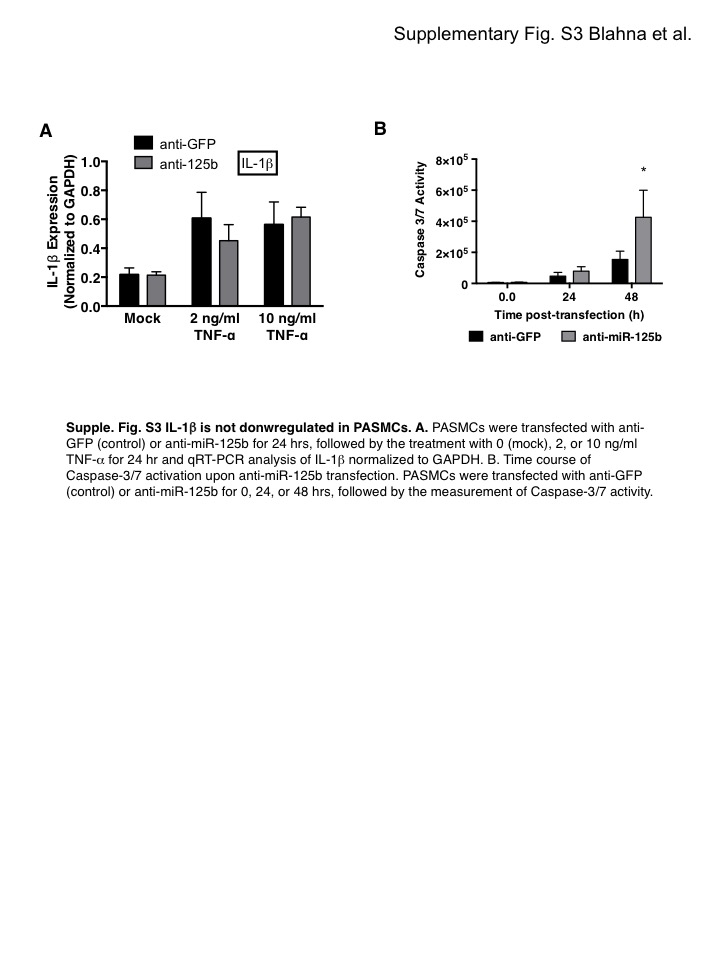

### Supplemental Fig. S4

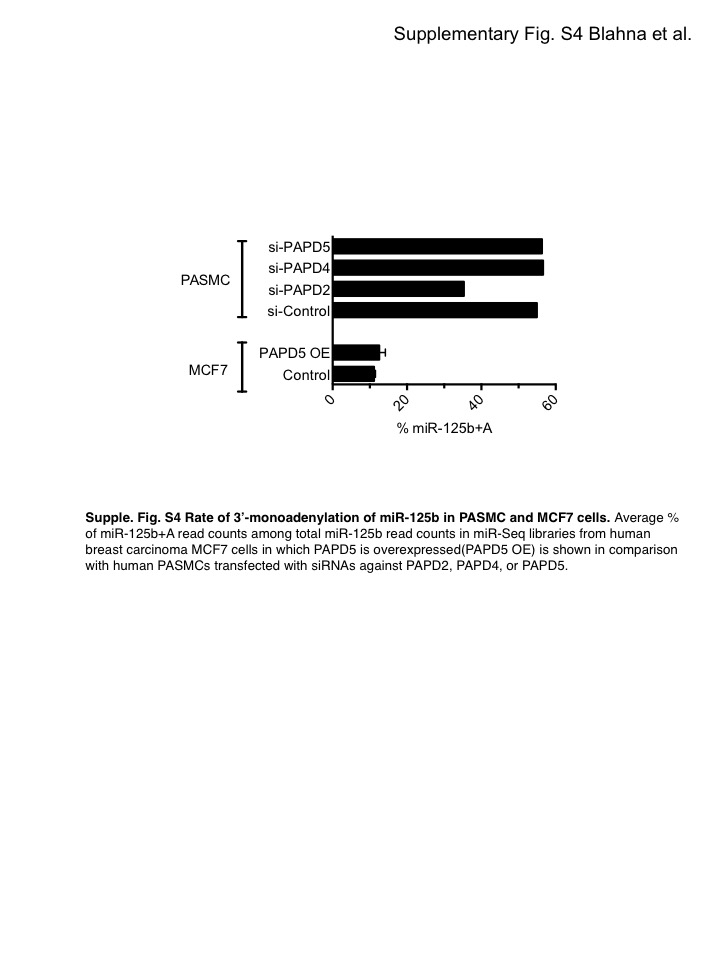

### Supplemental Fig. S5

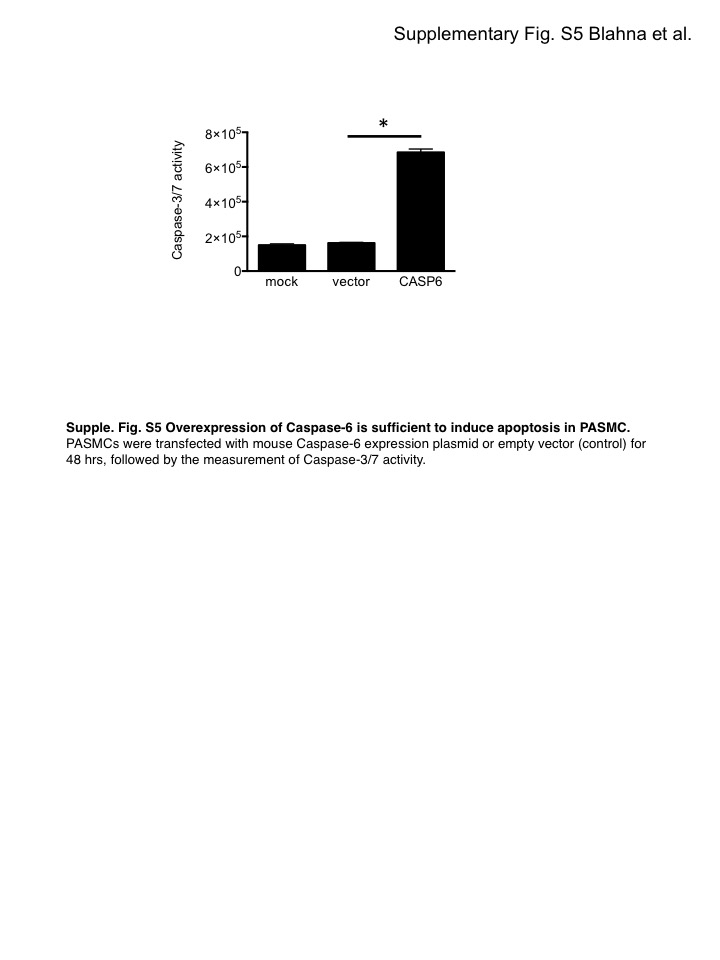

### Supplemental Fig. S6

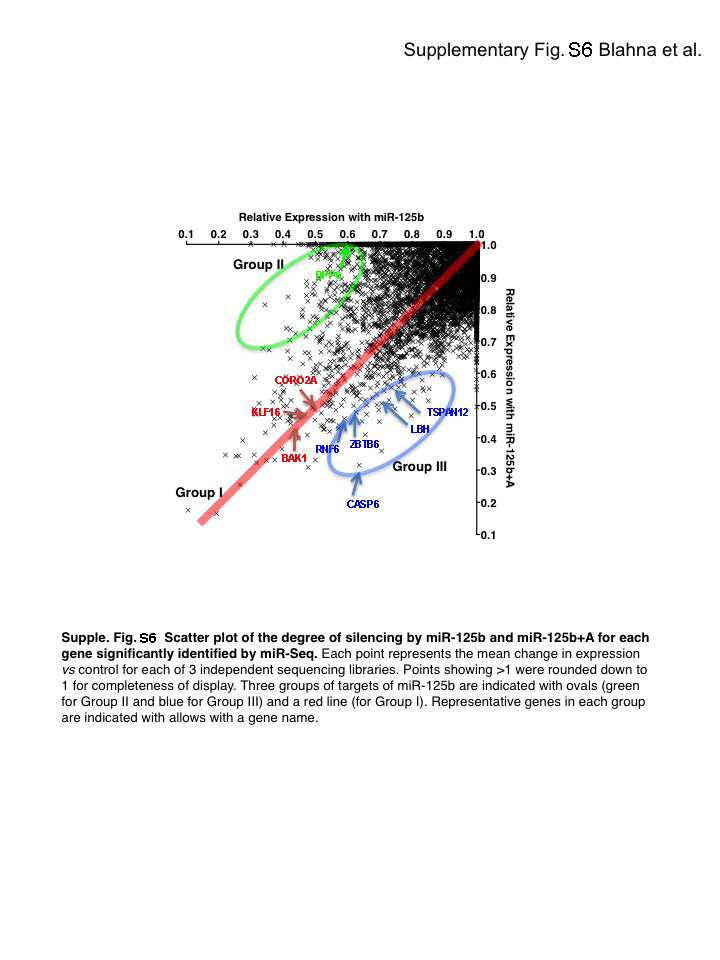
